# DMT1 contributes to MF- 438 - mediated inhibition of glioma cells

**DOI:** 10.1101/2024.04.26.591407

**Authors:** Laurence Zhang, Tomohiro Yamasaki, Tyrone Dowdy, Mioara Larion

## Abstract

Elevated SCD1 expression has been associated with enhanced cancer cell survival, proliferation, and resistance to therapy in many cancer types including gliomas. Hereby, we investigate the impact of MF-438 on SCD1-mediated lipid metabolism and its consequences on glioma growth and survival. Our data reveals an IDH^mut^-specific inhibitory effect of MF438 on gliomas. Also, we delineate a dual mechanism of action: while SCD1-mediated lipid metabolism is hindered by MF-438 treatment, MF-438 also exerts an SCD1-independent inhibition on DMT1 expression. Supporting data from the DMT1 blocker underscores its significance in MF-438’s anti-glioma efficacy.

## Introduction

Glioma, characterized by its aggressive nature and resistance to conventional therapies, remains a significant challenge in clinical. The pursuit of novel therapeutic strategies is imperative for improving patient outcomes in this disease. Recent studies have identified stearoyl-CoA desaturase 1 (SCD1) as a key enzyme in lipid metabolism that is implicated in the development and progression of various cancers, including glioma [1-3]. SCD1 catalyzes the synthesis of monounsaturated fatty acids (MUFAs) from saturated fatty acids, thereby playing a pivotal role in maintaining cellular lipid homeostasis. Interestingly, elevated SCD1 expression has been associated with enhanced cancer cell survival, proliferation, and resistance to therapy [4]. These findings have sparked interest in targeting SCD1 as a potential therapeutic strategy to overcome cancer resistance to treatment.

In this context, MF-438, a selective inhibitor of SCD1, emerges as a compelling therapeutic agent, including for gliomas. By specifically targeting SCD1, MF-438 has been demonstrated to reduce cell viability through the regulation of lipid metabolism and ER stress [5, 6]. Beyond these, MF-438 is also noted for its ability to modulate cancer stemness through multiple approaches [7-9]. A recent study revealed the potential of MF-438 in increasing the radiation sensitivity of esophageal squamous cell carcinoma cell lines, thereby resulting in an enhanced immune ferroptosis [10]. Furthermore, the sensitivity of MF-438 in glioma cell lines is associated with SCD expression. However, the mechanism by which MF-438 contributes to anti-glioma effects remains underexplored. Hereby, we investigate the impact of MF-438 on SCD1-mediated lipid metabolism and its consequences on glioma growth and survival. Our data reveals an IDH^mut^-specific inhibitory effect of MF438 on gliomas. Also we delineate a dual mechanism of action: while SCD1-mediated lipid metabolism is hindered by MF-438 treatment, MF-438 also exerts an SCD1-independent inhibition on DMT1 expression. Supporting data from DMT1 blocker underscores its significance in MF-438’s anti-glioma efficacy.

## Materials and Methods

### Cell Culture

Glioma cell lines: TS603 and GSC923, were cultured with DMEM: F12 media (Gibco Laboratories, Gaithersburg, MD, USA) with the supplement of penicillin/streptomycin, N2 growth supplement, epidermal growth factor (EGF) and fibroblast growth factors (FGF) (ThermoFisher Scientific, Waltham, MA, USA). U251_R132H and U251_IDH were cultured with DMEM with the supplement of 10%FBS.

### Metabolite Extraction

Cell pellets were collected at 48 hrs post-treatments and sonication by Misonix XL-2000 Ultra-liquid processor (Misonix Inc., Farmingdale, NY, USA) at 40 amps for 30 s. Following sonication, metabolite extraction was prepared using Methanol:Chloroform:water protocol and stored at -80 °C as previously described [11].

### Cell Proliferation Assay (CCK-8)

15000 cells per well (Ts603 and GSC923) or 2000 cells (U251_R132H and U251_WT) per well were seeded at the round bottom 96-well plate for six repeats. After the treatment of drugs, the cells were continued cultured in a cell CO_2_ incubator at 37 °C. CCK8 reagent (10ul/well) was added into the cells at 72hrs post-treatment and the absorbance at 450 nm was examined using a microplate reader.

### TBARS assay

0.5 x 10^6^ cells (Ts603 and GSC923) or 0.1 x 10^6^ cells (U251_R132H and U251_WT) per well were treated with MF-438 for 3 days. The cells were collected at indicated time and lysis with cell lysis buffer 3 (Cat No: 895366). After acid treatment, the assay was followed instructions (R&D, Cat No: KGE013). The results were examined and read at 450 nm using a microplate reader. The protein concentration of each sample was used to normalize the data.

### Colony formation assay

0.5x10^5^ Ts603 or GSC923 were cultured in 0.5% low-melt temperature agarose gel. The cells were treated with 1uM MF-438 or DMSO or 4.3 μM DMT1 blocker2 every 2∼3 days. The number and diameter of the colony were counted using the Celigo machine at day 0 and day 21.

### Western Blot

The cells were collected and lysis using SDS lysis buffer. Equal amounts of protein were loaded in each lane for 4-15% Bio-Rad Mini Protean TGX gel and then transferred to polyvinylidene difluoride membranes. The membranes were washed with blotting buffer (1× PBS containing 0.1% Tween20) and then blocked for 60 min in blotting buffer containing 10% low-fat powdered milk. Membranes were washed 3 times with blotting buffer, incubated at 4° C overnight with primary antibody (1:1000) containing 5% lowfat powdered milk, and incubated with secondary antibody (1:1000) at room temperature for 60 min. Target proteins were developed with Bio-Rad Clarity Western ECL substrate. GPX4 (Cat No. 52455), DMT I (Cat No. 15083), FTHI (Cat No. 4293S), beta-actin (Cat No. 8547), was purchased from cell signaling tech Inc. . SCD1 was purchased from Abcam (Cat No. ab19862). The blots were detected with BioRad ChemiDoc Touch imaging system. The relative expression of proteins was normalized to beta-actin and analyzed using Image J.

## Results

### MF-438 treatment results in a slow-down cell growth of glioma cells in vitro

To elucidate the underlying mechanism of MF438 against glioma stem cells, we treated IDH1^mut^ oligodendroglioma (TS603) and IDH1^WT^ GBM (GSC923) cell lines with various doses of MF-438 for three days. As depicted in Figure 1A, MF438 exhibited a dose-dependent inhibitory effect on TS603 cells with a dose of 1μM achieving a significant reduction of glioma cell growth. However, the efficacy of MF438 in inhibiting GSC923 growth was limited, demonstrating a challenge to achieve an IC50 reduction in cell proliferation. A more pronounced inhibition was observed in highly overexpressing IDH1^mut^ model system of the glioblastoma cell line (U251 R132H; U251^RH^) compared to IDH^WT^ overexpressing counterparts (U251^WT^) cells (Figure 1B). Nevertheless, a higher dose of MF-438 (10 μM) was required for the U251^WT^ cell lines (Figure 1A-1B). To exclude potential off-targets of MF438, we assessed the influence of SCD know-down on the cell grown. A similar reduction of cell growth was observed in both U251^RH^ and U251^WT^ cells 48 hrs post-siRNA SCD treatment (Figure 1C). Moreover, the addition of saturated fatty acids (SFA) increased the effect of MF-438 on the viability of TS603. (Figure 1D). To validate the drug’s effect, a colony formation assay was conducted to assess the long-term cell survival following 3 weeks of MF-438 treatment (Figure 2A). The results indicate a significant decrease in the number of colonies in MF-438-treated TS603 and GSC923 cells. Additionally, neutrospheres from U251^RH^ and U251^WT^ were cultured with or without 10 μM MF438. Limited dilution analysis was used to measure the sphere formation frequency, which was lower in MF-438 treated U251^WT^ and U251^RH^ cells compared to controls. Collectively, these data confirm the inhibitory effects of MF-438 on glioma in vitro and suggest a specific effect related to IDH1 mutation, since these cells are more sensitive to MF-438 inhibition.

**Figure 1.**
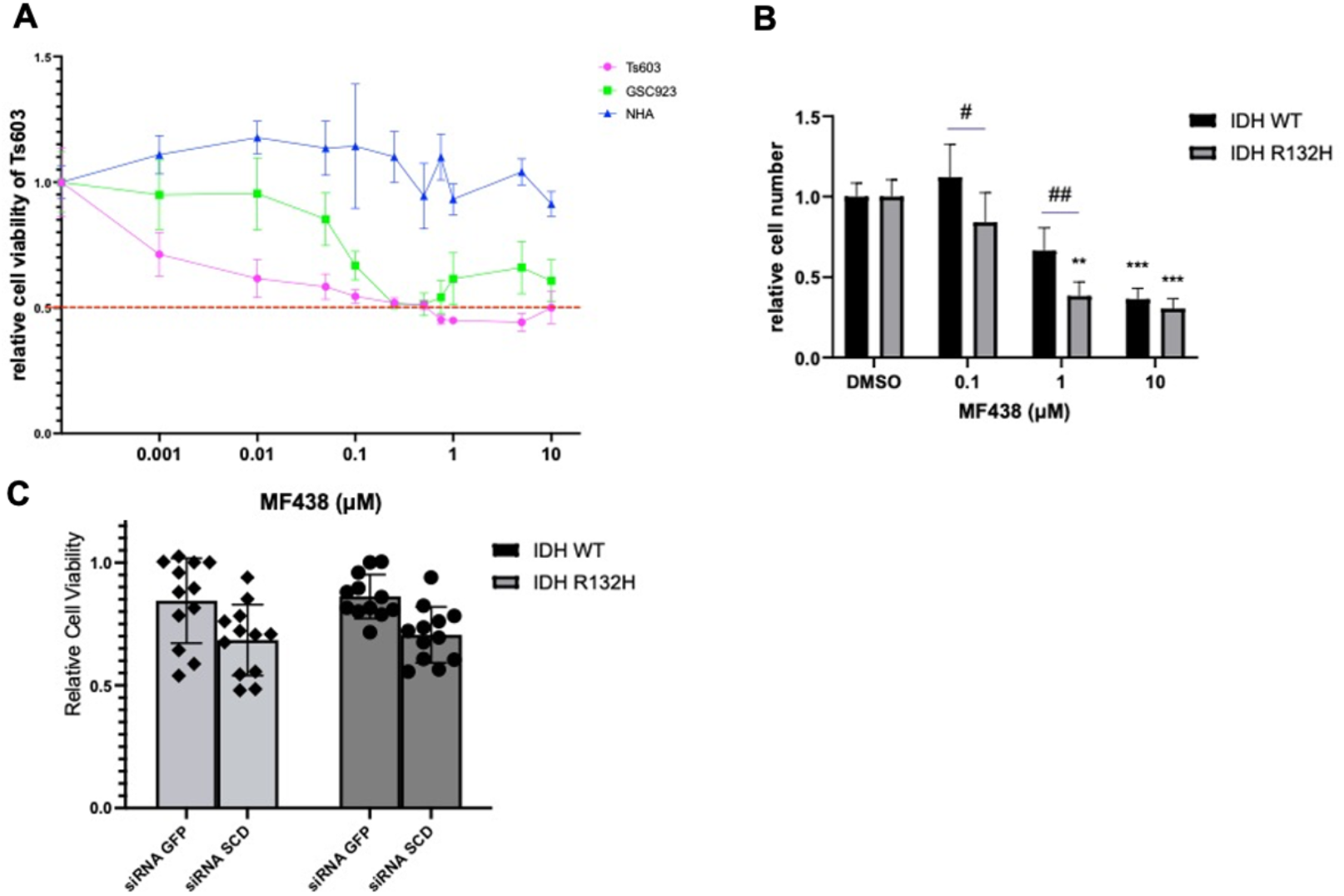
MF-438 inhibits the glioma cell growth. (A) EC50 measurement was performed to evaluate the effects of MF-438 on the cell growth of two patient-derived glioma cell lines: TS603, GSC92. NHA was used as a control (N=6). (B) The cellular viability of U251 RH and U251WT were examined by CCK-8 assay after MF-438 treatment. (C) The impact of SCD knockdown on cell growth was examined at 48h post-siRNA treatment. (D) The impact of MF-438 on SFA stimulation was evaluated using CCK-8 assay (N=6).

**Figure 2:**
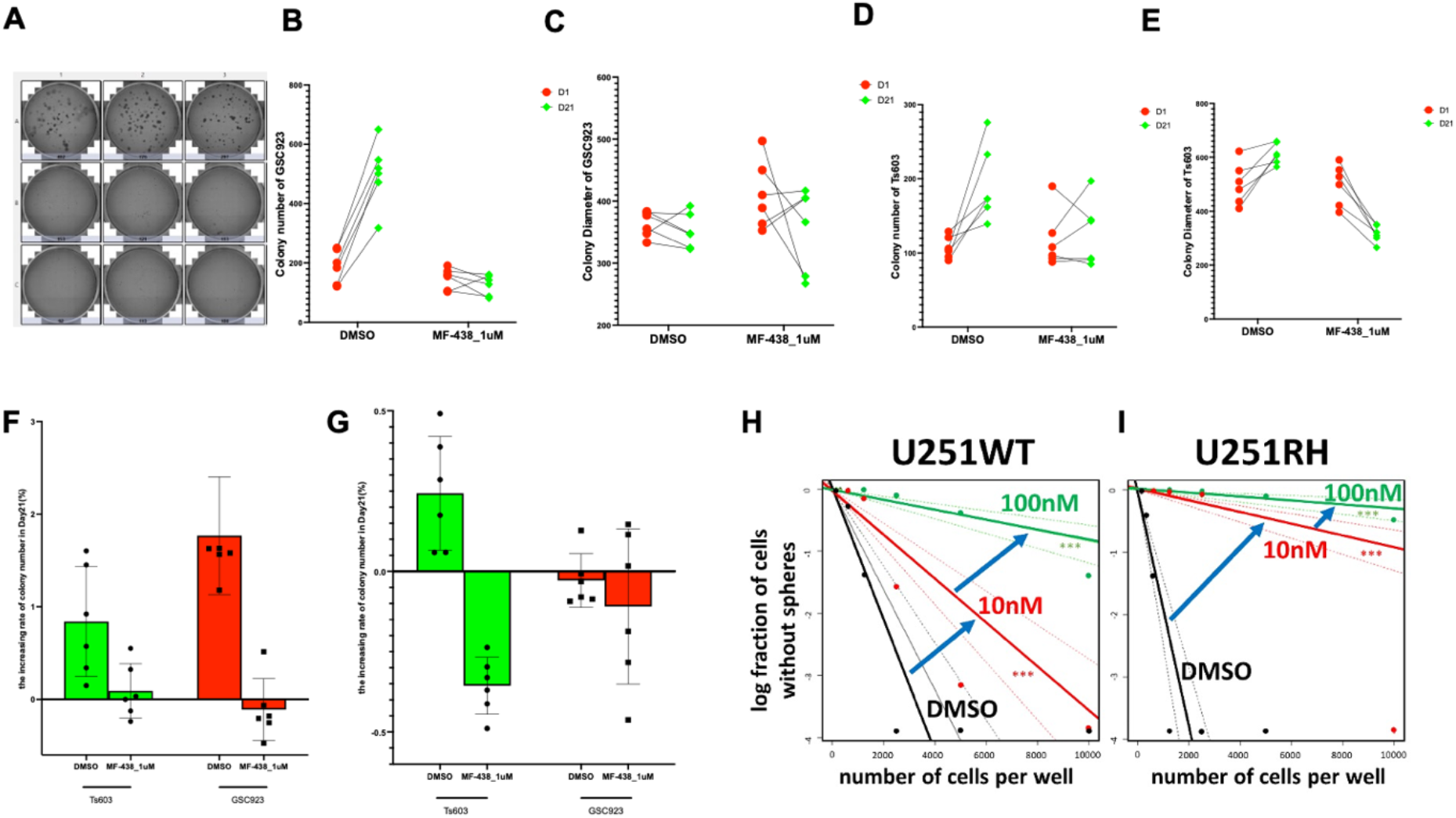
MF-438 treatment reduces the colony formation in vitro. (A-G) 0.5x10^5^ Ts603 or GSC923 were cultured in 0.5% low-melt temperature agarose gel. The impact of MF-438 on cell survival was examined using colony-forming (CFU) assay. (H and I) A limited dilution assay was used to influence MF-438 on neutrosphere formation of U251RH and U251WT.

### MF-438 decreases SCD-mediated FA metabolism

To explore the impact of MF-438 treatment on fatty acid metabolism, the cell growth was examined with or without SFA stimulation. No significant effect on glioma proliferation was observed with different doses of SFA, even after repeated SFA stimulation (Figure 3A). Interestingly, SFA stimulation had no impact on cell proliferation in SCD-silenced glioma cells compared to BSA control, suggesting that SFA-induced FA metabolism does not contribute to cell survival despite being a major substrate of the SCD enzyme (Figure 3B-3C). Surprisingly, SFA stimulation produced a synergistic inhibitory effect on glioma growth when combined with 1 μM MF-438 treatment, indicating that SFA might have an SCD-independent effect that was added to MF-438-mediated effects (Figure 3D).

**Figure 3.**
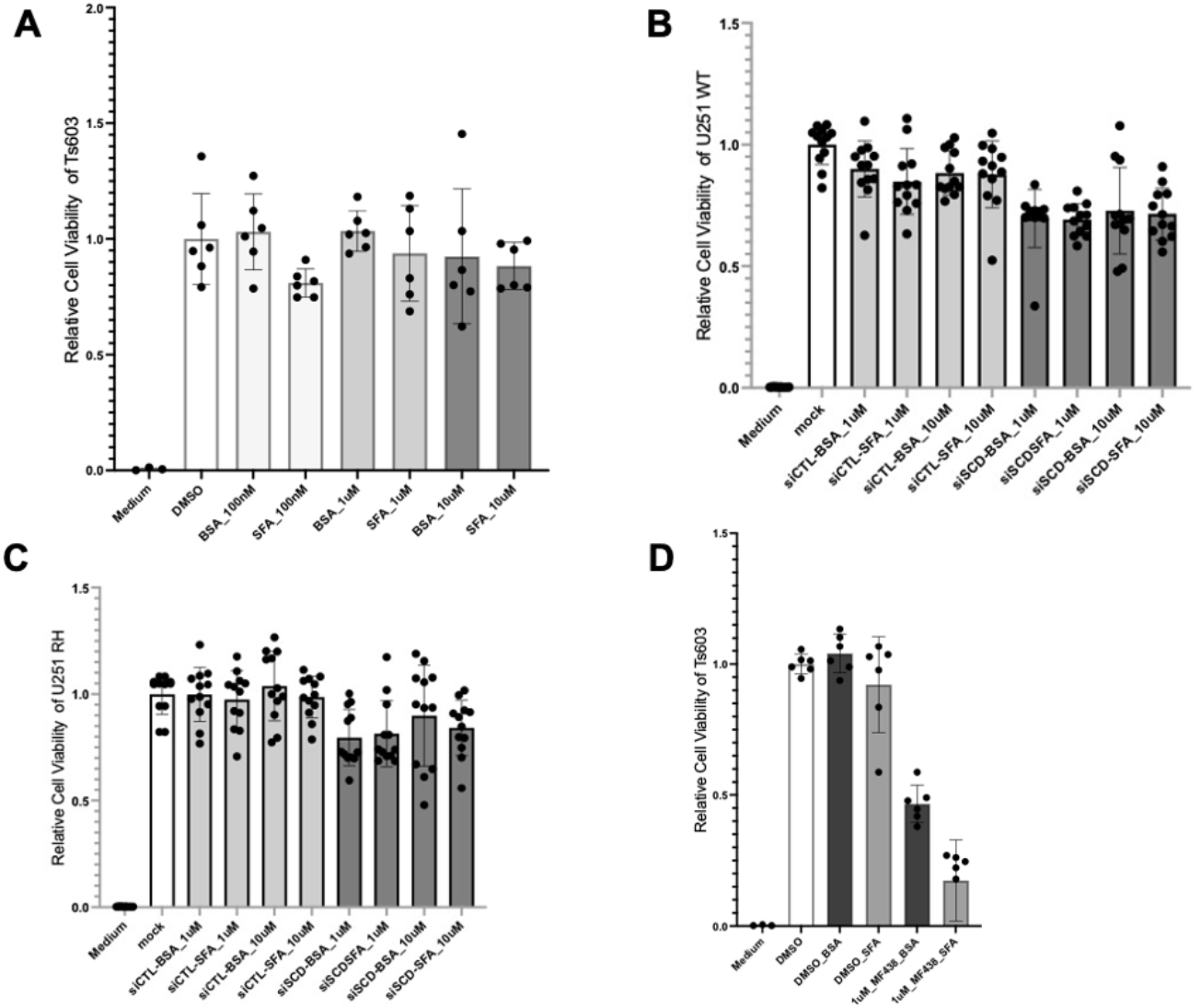
SFA stimulation has no impact on SCD silencing mediated cell growth. (A) Ts603 cells were stimulated with different concentration of SFA or BSA. The cell growth was monitored with cck8 assay (N=6). (B and C) U251 WT(B) and U251(C) were treated with siRNA-SCD1 or siRNA-GFP for 48hrs. Then the cells were stimulated with 10uM SFA or BSA for 48hrs. The cell growth was examined with cck8 asay.

To better understand the metabolic response to MF-438-mediated SCD inhibition, a targeted lipidomic analysis using LC/MS evaluated the lipid species in glioma cells before and after MF438 treatment in TS603 and GSC923 cell lines. As shown in Figure 4, the levels of monounsaturated fatty acids MUFAs (C16:1 an C18:1) were significantly reduced by inhibition of SCD, with MF-438 more pronounced in TS603 (Figure 4A-4B) than GSC923 (Figure 4C-4D). Interestingly, 10nM MF-438 treatment resulted also in a decrease in the abundance of SFAs in TS603 cells, which is unexpected since SFA would go up if the SCD gets inhibited. This suggests that cells shunt SFAs into other pathways to avoid cytotoxicity.

**Figure 4.**
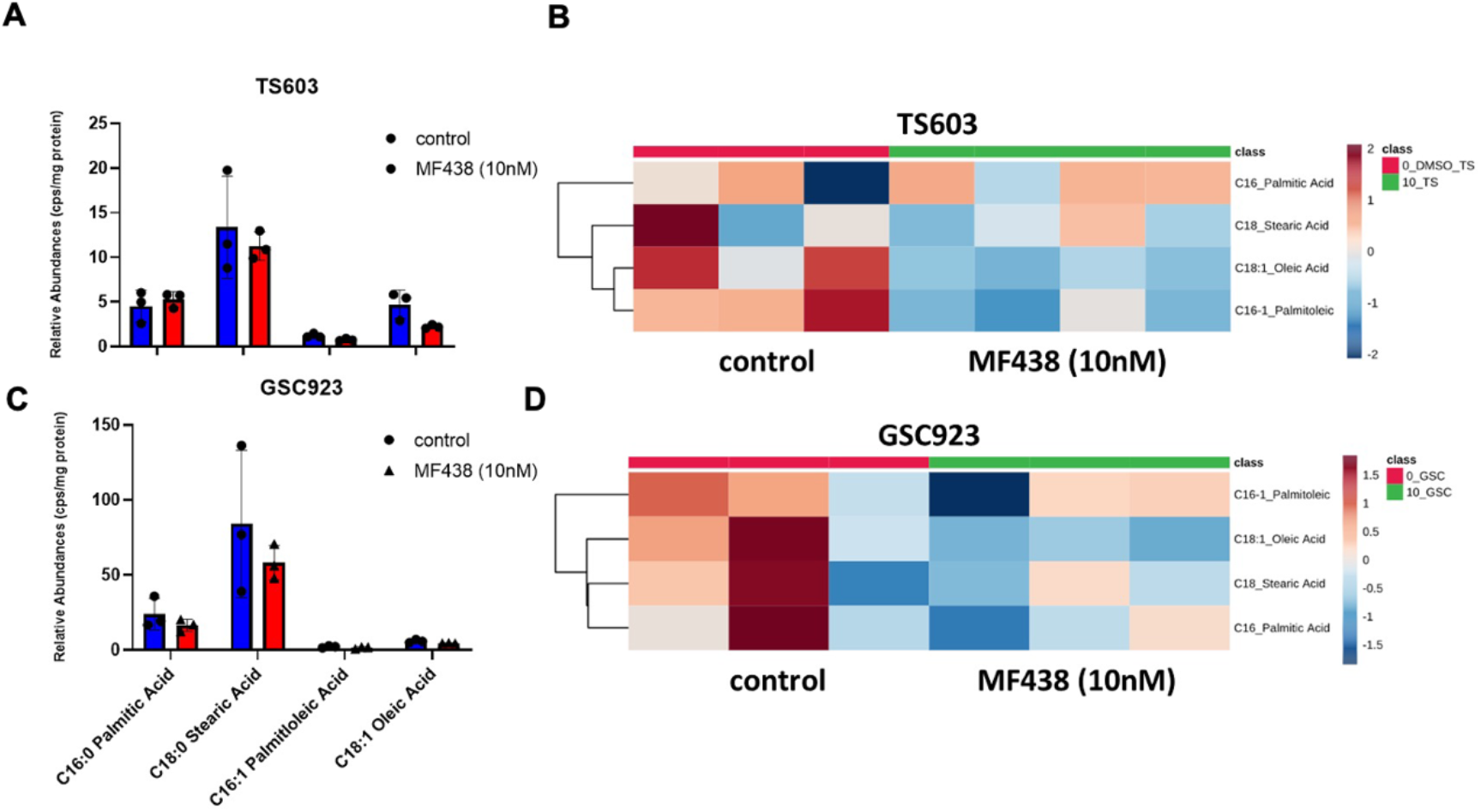
MF-438 treatment reduces the abundance of MF438 in Ts603 cells. LC/MS was performed to measure the abundance of SFA and MUFA in Ts603 (A), GSC923 cells (B), U251 WT(C) and U251 RH(D).

### Ferroptosis contributes to MF-438-mediated cell death

Given that MUFA has been implicated in a ferroptosis resistance status in cancer cells, we investigated whether MF-438 mediated MUFA reduction could induce a concomitant ferroptosis in glioma cells. Examining the ratio of NAPD+/NADPH and PUFA peroxidation (Figure 5A and 5B) revealed that MF-438 treatment increases PUFA peroxidation in TS603 cells, but not GSC923 cells, indicating a following ferroptosis induced by MF-438 treatment in TS603 cells. As expected, GPX4, a ferroptosis marker, was significantly decreased by MF438 treatment in Ts603 cells. Other ferroptosis related markers, FTH1 and DMT1, were also downregulated in MF-438-treated glioma cells (Figure 6A). Although no decrease in GPX4 expression was observed in U251 RH cells with MF-438 treatment, the expression of FTH1 and DMT1 exhibited a similar reduced pattern as in TS603 cells. However, SCD silencing had no impact on the expression of DMT1 and FTH1 (Figure 6B). More importantly, the reduction of DMT1 by MF-438 was dose-dependent, suggesting that DMT1 plays a key role to MF438 induced ferroptosis in glioma cells, independent of SCD contribution (Figure 6C).

**Figure 5.**
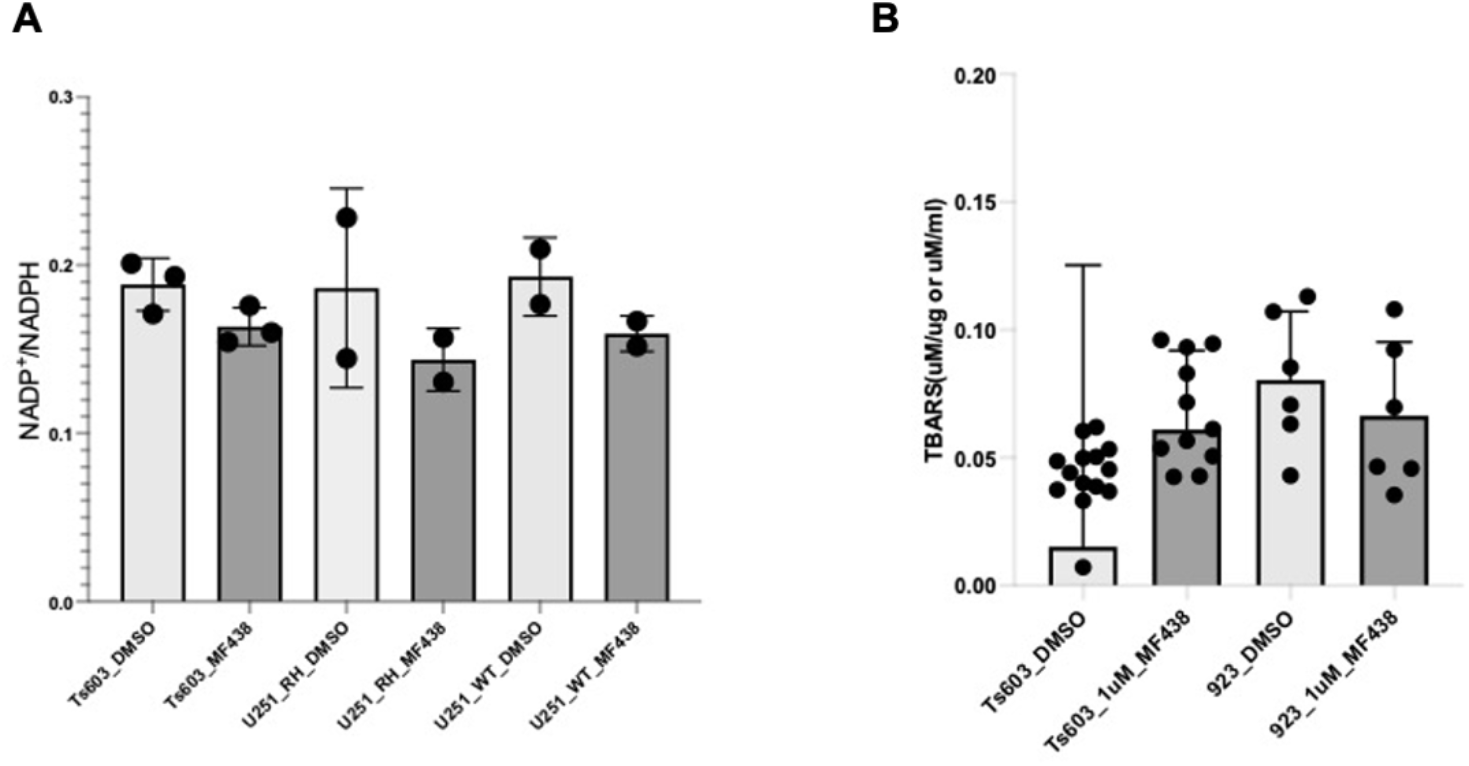
MF-428 treatment increases lipid peroxidation in Ts603 cells. A) The cells were treated with MF-438 for 3days. The cell lysates were collected. NADP+/NADPH were measured and normalized with protein concentration. B) lipid peroxidation was examined using TBARS assay after 3 days of MF438 treatment. The data was normalized with protein concentration.

**Figure 6.**
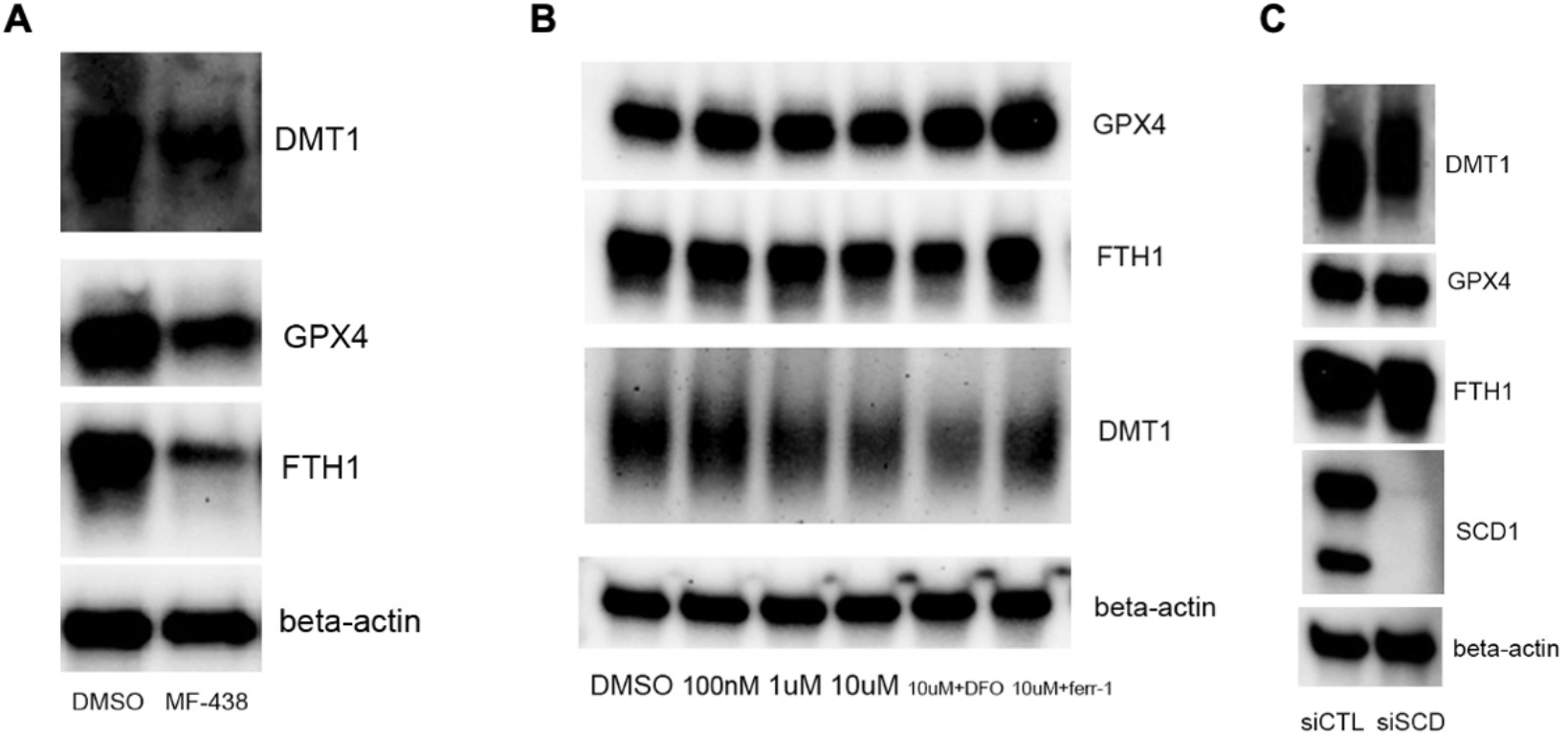
Ferroptosis is induced in Ts603 cells. A) DMT1, GPX4 and FTH1 expression were examined using western blotting after 3 days treatment of MF-438 in Ts603 cells. B) DMT1, GPX4 and FTH1 expression were examined using western blotting after 3 days of treatment using different concentrations of MF-438 in U251 RH cells. ( C ) DMT1, GPX4, FTH1 and SCD expression were examined using western blotting after siRNA-SCD treatment in U251 RH cells.

To further address the role of DMT1 in glioma, we examined its expression across different glioma cell lines. Interestingly, DMT1 expression was detected in most glioma cell lines, but not in normal human astrocytes (NHA) cell lines, even with overexpression of IDH1 R132H in NHA cells (Figure 7A). Blockage of DMT1 in glioma cells significantly decreased cell viability 48 hours post-treatment. Colony forming (CFU) assay showed that the DMT1 inhibitor significantly reduced the number of colonies formed (Figure 7B∼D). Collectively, these data suggest that MF-438 inhibits the glioma cells’ growth partly through the downregulation of DMT1.

**Figure 7.**
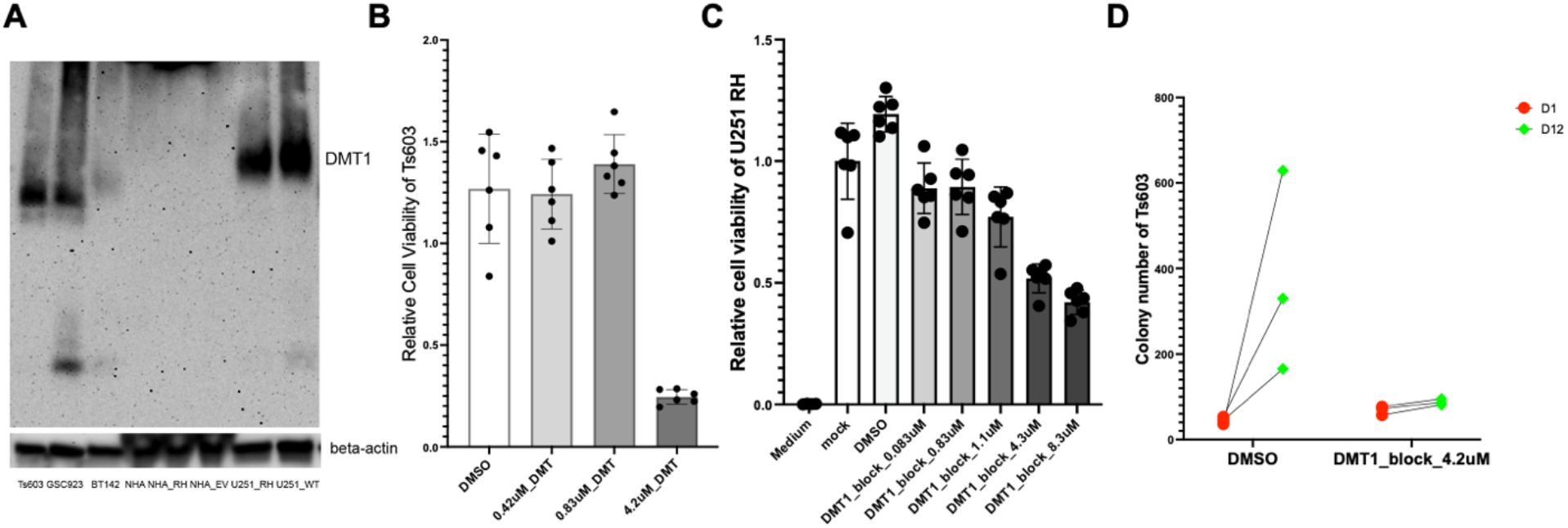
DTM1 treatment significantly reduces the cell viability of IDH mutation glioma cells. A) DMT1 expression in different glioma cell lines was examined using western blotting. B and C) The impact of DMT1 blocker on Ts603 (B) and U251 RH (C) was examined using cck8 assay (D) CFU assay was used to evaluate the long-term effect of DMT1 blocker on Ts603 survival.

## Discussion

In this study, our data demonstrate that MF-438 treatment inhibits the growth of glioma cells. This treatment disrupts a positive feedback loop between SFA and MUFA. A reduction of intracellular MUFA and increased polyunsaturated fatty acids (PUFAs) peroxidation are suggested to be associated with ferroptosis in IDH1 mutation glioma cells, supported by the up-regulation of ferroptosis markers following MF-438 treatment.

The high content of SFA in cell membranes and increased intracellular MUFA production have been identified as major factors contributing to the chemotherapeutic resistance of cancer cells. A recent study by Magtanong et al., suggests that exogenous MUFAs can prevent cell ferroptosis by blocking the accumulation of lipid ROS and reducing the levels of PUFA-phospholipids in the cell membrane in an ACSL3-dependent manner [12]. Supporting this, breast cancer cells supplemented with MUFAs have been shown to be resistant to combination treatment with ferroptosis inducers: CCI-779 and RSL3. More importantly, the crosstalk between SCD and SREBP-1 was found to be crucial for establishing ferroptosis resistance. Knockout of SCD or the use of SCD1 inhibitors rendered the cancer cells more sensitive to ferroptosis inducers, whereas, SCD1 overexpression did not [13]. As a key enzyme of fatty acid synthesis *de novo*, SCD is responsible for the insertion of the first double bond into the Δ9 position of fatty acyl-CoA substrates to generate MUFAs [4]. SCD silencing has been shown to reduce MUFA levels and increase fatty acid oxidation through upregulation of CTP1 activity. Furthermore, the increased ratio of SFA and MUFAs is considered as a metabolic marker of lipotoxicity. High concentrations of palmitic acid (>25M) have increased lipo-apoptosis in SCD-deficient cells [14]. SFA (16:0) has been shown to enhance the anti-proliferation effects of SCD inhibitors on cancer cells [15]. In our study, although SCD silencing suppressed glioma cell growth, a low dose of SFA had no significant impact on cellular death. Moreover, contrary to expectations, SFA 16:0 was significantly reduced instead of increased. The abundance of MUFAs was also significantly decreased in cells treated with siRNA SCD and MUFA-treated IDH^mut^ cells, indicating MF-438 treatment may counteract MUFA-enforced ferroptosis resistance. In this context, ASCSL3 expression was slightly decreased. Most interestingly, we did not observe an induction of GPX4 in either MF-438-treated or siRNA SCD-treated glioma cells. However, DMT1 expression was significantly affected by MF-438 treatment, exhibiting a dose-dependent inhibition and suggesting that iron metabolism is reprogrammed by MF-438. Iron-mediated PUFAs peroxidation may contribute to MF-438-mediated ferroptosis. As predicated, MF-438 treatment increased PUFAs peroxidation products in TS603 cells. Moreover, siRNA SCD did not reduce the expression of FTH1 and DTM1, indicating a MF-438-specific effect on cell death.

In brain, DMT1 mainly distributes in neuronal cells [16], but is expressed at low levels in astrocytes and oligodendrocytes. However, a comparison of DMT1 among glioma cell lines revealed prominent expression in these cell lines, but not in NHA-derived cell lines, indicating that DMT1-dependent iron uptake plays an important role in the iron metabolism of glioma cells. Interestingly, a recent study showed that a high dose of Temozolomide(200uM) resulted in ferroptosis through upregulation of DMT1 in TG905 cells [17]. Given the evidence of a reverse effect on TMZ-mediated inhibition by DMT1 silencing, we investigated the contribution of DMT1 inhibition to the proliferation of glioma cells. Inspiringly, a DMT1 blocker produced a profound inhibitory effect on the growth of glioma cells.

In conclusion, our *in vitro* data indicate that MF-438 treatment induces ferroptosis and suppresses the growth of glioma cells. This inhibitory effect is partly dependent on the reduction of DMT1 following treatment. More importantly, our data shows that DMT1 blocking significantly induces the death of glioma cells by itself, providing a novel strategy to augment the metabolic vulnerabilities of glioma.

## Author Contributions

Conceptualization, L.Z. and M.L.; methodology, L.Z. T.D., T.Y.., and M.L; formal analysis, L.Z T.D, and M.L.; data curation, T.D. and L.Z; writing—original draft preparation, L.Z.; writing—review and editing, M.L.; supervision, M.L.; project administration, M.L.; funding acquisition, All authors have read and agreed to the published version of the manuscript.

## Funding

This research was funded by the Intramural Program of the National Cancer Institute. This research was supported by the National Institutes of Health Intramural Research Program through an NCI FLEX award to A.L. M.R.G. and M.L. entitled “Live cell metabolism via Raman imaging microscopy.”

### Acknowledgments

We would like to thank Dr. ChunZhang Yang (NOB) for providing us with the U251^*WT*^ and U251 ^*R132H*^ constructs and Dr. Timothy Chan for providing us with TS603 cells.

